# Multifaceted Roles of Histone Lysine Lactylation in Meiotic Gene Dynamics and Recombination

**DOI:** 10.1101/2024.01.25.576681

**Authors:** Xiaoyu Zhang, Yan Liu, Ning Wang

## Abstract

Male germ cells, which are responsible for producing millions of genetically diverse sperm through meiosis in the testis, rely on lactate as their central energy metabolite. Recent study has revealed that lactate induces epigenetic modification in cells through histone lactylation, a post-translational modification involving the addition of lactyl groups to lysine residues on histones. Here we report dynamic histone lactylation at histone H4-lysine 5 (K5), -K8, and -K12 during meiosis prophase I in mouse spermatogenesis. By profiling genome-wide occupancy of histone H4-K8 lactylation (H4K8la), which peaks at zygotene, our data show that H4K8la mark is observed at the promoters of genes exhibiting active expression with Gene Ontology (GO) functions enriched for meiosis. Notably, our data also demonstrate that H4K8la is closely associated with recombination hotspots, where machinery involved in the processing DNA double-stranded breaks (DSBs), such as SPO11, DMC1, RAD51, and RPA2, is engaged. In addition, H4K8la was also detected at the meiosis-specific cohesion sites (marked by RAD21L and REC8) flanking the recombination hotspots. Collectively, our findings suggest that histone lactylation serves as a novel mechanism through which lactate regulates germ cell meiosis.

## Introduction

Meiosis is a specialized form of cell division with the primary aim of producing haploid gametes from diploid germ cells in sexually reproducing organisms (*1*). To occur, the activation of genes crucial for meiosis plays a pivotal role in expressing the molecular machinery responsible for implementing meiotic homologous recombination (*2*). The initiation of meiotic recombination involves the programmed creation of DNA double-strand breaks (DSBs) facilitated by the conserved type II topoisomerase-like enzyme SPO11. A specific subset of these meiotic DSBs undergo resolution as crossovers, entailing the reciprocal exchange of DNA between homologous chromosomes. It is noteworthy that these DSBs are not randomly distributed along eukaryotic chromosomes; instead, they preferentially form in permissive regions referred to as hotspots.

The epigenetic regulation of meiotic recombination hotspots involves various molecular mechanisms. For examples, DNA methylation establishes domains of meiotic recombination along chromosomes by silencing euchromatic crossover hotspots (*3*). Spatially accessible genomic regions during meiotic prophase are associated with DSB-favored loci (*4*). Perhaps the best characterized epigenetic regulation of meiotic recombination hotspots is post-translational modifications of histones, such as histone acetylation and methylation. Specific histone marks, such as H3K4me3 (trimethylation of histone H3 at lysine 4) and H3K36me3, are often enriched at active recombination hotspots.

The histone modification H3K4me3 frequently occurs at the transcription start site of genes actively expressed in eukaryotic cells. Given its correlation with gene expression levels, H3K4me3 is commonly thought to play an instructive role in gene expression, making it a histone modification with an ’activating’ function. In addition to its role in gene transcription, H3K4me3 is also known as a marker for hotspots DSB sites at the very earliest stages of meiosis (*5, 6*). In mammals, this is performed by the protein PRDM9, which is expressed in the leptotene and zygotene substages (*7, 8*). DSB formation at sites defined by PRDM9 is catalyzed by the SPO11 enzyme and its binding partner TOPOVIBL (*9–13*). SPO11-mediated cleavage results in single-strand DNA overhangs that are subsequently coated by various proteins, including DMC1 and RAD51 (*14–16*). The DSBs enable homology searching and alignment to occur, which in turn promote homology synapsis and DSB repair (*17*).

It has long been recognized that, different from most somatic cells, germ cells use lactate as their central energy metabolite (*18, 19*). Inside the seminiferous tubules, meiotic germ cells stay behind the blood-testis barrier. Sertoli cells, positioned as custodians of germ cell development, exhibit a unique metabolic profile favoring glycolysis and lactate production, which is released into the extracellular space. Monocarboxylate (MCT) transporters facilitate the transport of lactate, ensuring its efficient uptake by germ cells. Beyond its role as an energy substrate, lactate engages in signaling, influencing gene expression and participating in the intricate regulation of spermatogenesis.

Recently, in addition to its well-known metabolite function, lactate has recently been discovered to induce epigenetic modification in cells through histone lactylation. Histone lactylation is a post-translational modification involving the addition of lactyl groups to lysine residues on histones (*20*). Here, by mapping the genome-wide occupancy of histone H4 lysine 8 (K8) lactylation (H4K8la), our data reveal that histone lysine lactylation is associated with active gene expression, meiotic recombination hotspots, and the cohesion sites flanking these hotspots. These findings suggest novel roles of lactate in meiosis, with histone lactylation emerging as a key underlying mechanism in this intricate process.

## Results

### Dynamic histone lactylation during meiosis prophase I

We initially performed immunofluorescence staining on juvenile testicular cross sections to investigate various histone lysine lactylation marks. As a positive control, H3K4me3, known for its dynamic regulation during meiosis prophase I and essential roles in gene transcription and recombination, was used (*21*). Our data revealed dynamic lactylation of histone H4 at lysine 4 (K4), K8, and K12 during early meiosis (**Figure 1A**). To further characterize these histone H4 lysine lactylations, chromosome spreads were employed (**Figure 1B**). Our data indicate that the H4K5la mark persists from the leptotene to the pachytene stage. H4K8la exhibits a rapid increase from the leptotene stage, reaching its peak at the zygotene stage, and then quickly disappears at the pachytene stage. H4K12la is present at both leptotene and zygotene stages but disappears at the pachytene stage. Given the unique upregulation of H4K8la at the zygotene stage, we focused on H4K8la in this study.

**Figure 1.**
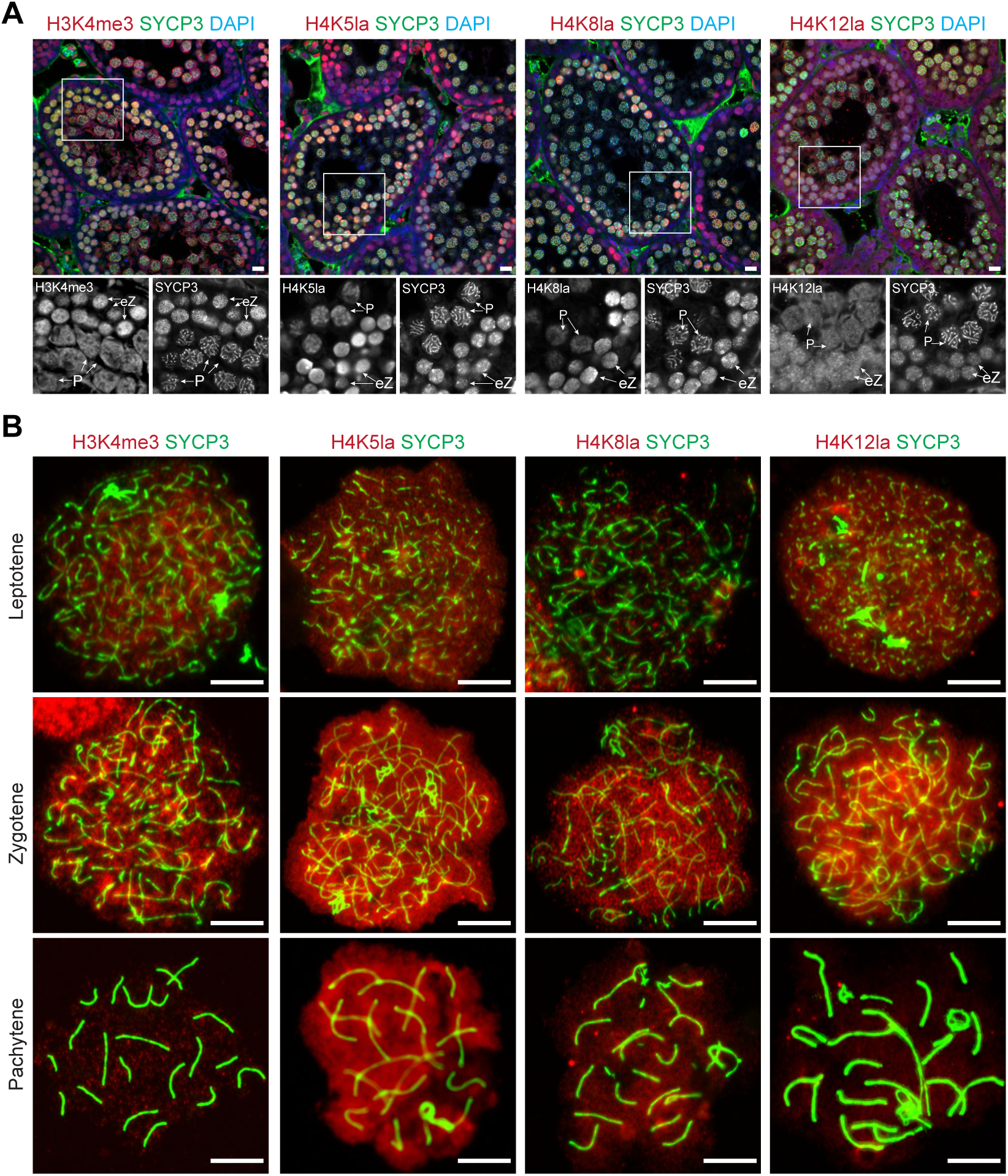
H3K4me3, H4K5la, -K8la and -K12la expression in meiotic germ cells. **(A)** Staining of mouse juvenile testicular cross-sections using antibodies against H3K4me3, H4K5la, H4K8la, H4K12la, and SYCP3. DAPI was used to stain nuclei. SYCP3 indicates meiotic germ cells. Scale bar: 20 μm. **(B)** Chromosome spread staining of H3K4me3, H4K5la, H4K8la, and H4K12la in germ cells at indicated meiotic stages. Scale bar: 20 μm.

### Characterization of genome-wide H4K8la occupancy

To enrich germ cells at the leptotene, zygotene, and pachytene stages of meiosis prophase I, we employed synchronized spermatogenesis (**Supplementary Figure 1**) (22). Subsequently, chromatin immunoprecipitation with sequencing (ChIP-seq) assays for H4K8la were performed using testicular cells enriched for leptotene, zygotene, and pachytene spermatocytes. The resulting reads were then mapped to the mouse genome (**Supplementary Figure 2**).

### H4K8la is associated with meiotic gene expression

First, we correlated H4K8la with dynamic transcriptomic changes during leptotene, zygotene and pachytene stages, which were profiled by single-cell RNA sequencing (scRNA-seq) of juvenile testes (**Supplementary Figure 3**). The characterization of leptotene, zygotene, and pachytene cells is defined by the testicular reference map (23). Pseudo-bulk analysis of differentially expressed genes (DEGs) from leptotene to pachytene showed three major clusters (**Figure 2A and 2B**). We then correlated DEGs with H4K8la occupancy at their promoters. Our results demonstrate a positive correlation between H4K8la occupancy at the transcription start sites (TSSs) of DEGs and their gene expression from leptotene to pachytene (**Figure 2C-2E**). Meanwhile, genes with highest H4K8la occupancy at leptotene are enriched for Gene Ontology (GO) functions related to mRNA processing, cell cycle phase transition, and chromosome segregation (**Figure 2F**). Genes with the highest H4K8la occupancy at zygotene are enriched for GO functions related to meiotic cell cycle and meiosis I (**Figure 2G**). Lastly, genes with the highest H4K8la occupancy at pachytene are enriched for GO functions related to cilium movement and fertilization (**Figure 2H**). Together, our data demonstrate that H4K8la is correlated with transcriptionally active chromatin, suggesting a role for H4K8la in facilitating gene expression in meiosis prophase I.

**Figure 2.**
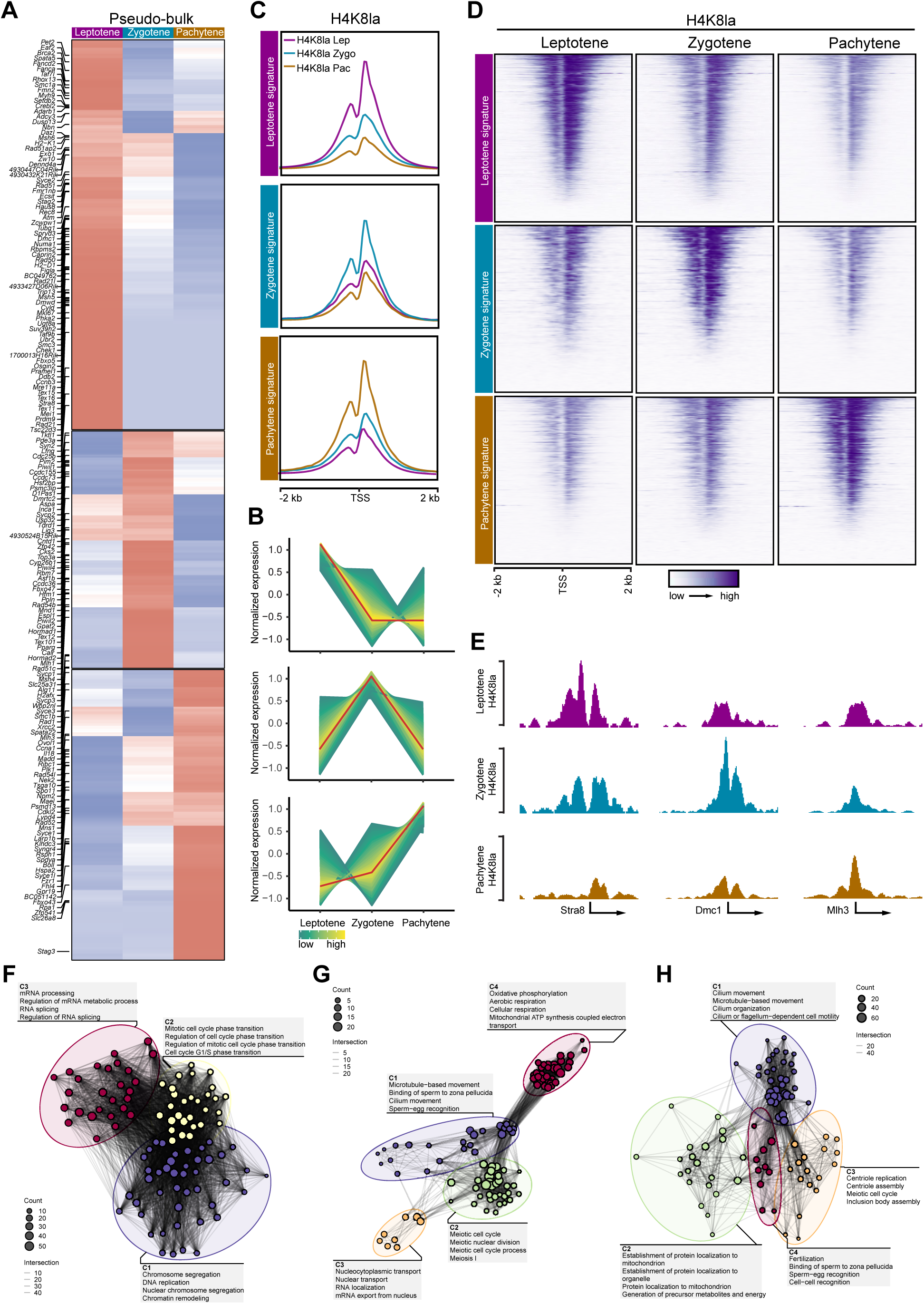
H4K8la dynamics at gene promoters from leptotene to pachytene at meiotic prophase I. **(A)** Pseudo-bulk heatmap of differentially expressed genes from leptotene to pachytene from a scRNA-seq dataset. Gene important for meiosis were labeled. **(B)** Stage-specific meiotic gene expression patterns. **(C)** Signal profiles for stage-specific H4K8la at TSSs recapitulate meiotic gene expression patterns. **(D)** Heatmap analysis for stage-specific H4K8la at TSSs recapitulate meiotic gene expression patterns. **(E)** Genome tracks showing H4K8la occupancy at the indicated gene promoter regions from leptotene to pachytene. **(F)** Network showing functional analysis of promoter regions with H4K8la peaks at leptotene. **(G)** Network showing functional analysis of promoter regions with H4K8la peaks at zygotene. **(H)** Network showing functional analysis of promoter regions with H4K8la peaks at pachytene.

### Dynamics H4K8la signals during meiotic recombination

Epigenetic marks play a crucial role in determine recombination hotspots (**Figure 3A**) (24). Given the similar pattern of H4K8la with H3K4me3 during meiosis prophase I, we wondered whether H4K8la marks are also present at the recombination hotspots. To identify these recombination hotspots, we conducted a comparative analysis of SPO11 oligos (SPO11-associated oligonucleotides) (*25*) and DMC1 SSDS (single-stranded DNA sequencing) (*26*). Our data show that the mean SPO11 oligo profile exhibits a narrow peak at the centers of hotspots, directly adjacent to the average DMC1-SSDS coverage (*27*) (**Figure 3B**; **Supplementary Figure 4A**). The DMC1-SSDS signal is confined to the DSB-initiating chromosome, reflecting the affinity of DMC1 to the 3′ ssDNA overhangs (**Figures 3C; Supplementary Figure 4B**).

**Figure 3.**
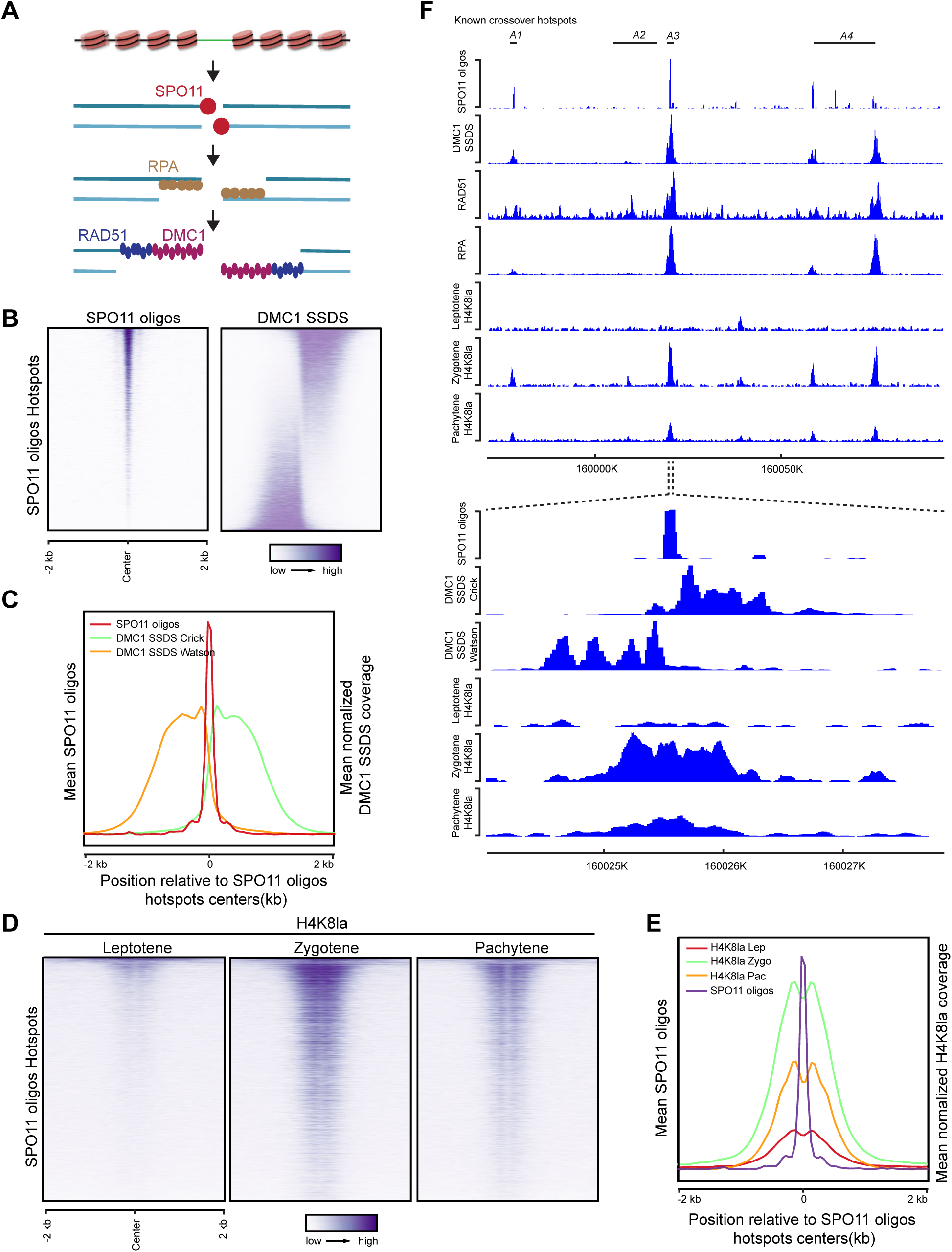
H4K8la dynamics at meiotic recombination hotspots. **(A)** A schematic for DSB formation and repair. Break sites are initiated by SPO11. This process is followed by a 5′-to-3′ strand resection, resulting in the formation of single-stranded DNA (ssDNA) overhangs. These overhangs are initially bound by RPA and subsequently bound by RAD51 and DMC1. **(B)** Heatmaps showing the coverage of SPO11 oligos and DMC1 SSDS in hotspots in the C57Bl/6 strain. Each row in these heatmaps corresponds to one of the hotspots. These hotspots are organized in descending order based on the abundance of SPO11 oligos. **(C)** Distribution of SPO11 oligos and both Crick and Watson strands of DMC1 SSDS coverage around the centers of SPO11 oligo hotspots. **(D)** Heatmaps showing the dynamic of H4K8la at SPO11 oligo hotspots. Each row in these heatmaps represents a specific hotspot-associated peak, extending ±2 kb from the peak’s center. These peaks are organized in descending order, based on their SPO11 oligo density. **(E)** Distribution of SPO11 oligos and hotspots associated with H4K8la peak coverages from leptotene to pachytene around the centers of SPO11 hotspots. **(F)** Genome browser view of SPO11 oligos, DMC1 SSDS, RAD51 ChIP-seq, RPA ChIP-seq, and H4K8la occupancy maps at four known crossover hotspots (*A1–A4*). SPO11 oligos, Crick- and Watson strands of DMC1 SSDS coverage and H4K8la occupancy in zoom-in window around hotspot *A3*.

Intriguingly, we observed a dynamic H4K8la signal at SPO11-oligo hotspots (**Figure 3D)** and DMC1-SSDS (**Supplementary Figure 4C**) hotspots from leptotene to pachytene: the H4K8la signal initially appeared at leptotene, reached its peak intensity at zygotene, and dissipated by pachytene. In both SPO11-oligo hotspots and DMC1-SSDS, the profile of H4K8la signal exhibit a significant enhancement in both the breadth and intensity of H4K8la marks during zygotene, followed by a decline at pachytene (**Figure 3E; Supplementary Figure 4D**).

We further conducted integrated analysis of H4K8la, SPO11-oligo, and DMC1-SSDS along with RAD51 and RPA2 sites, and analyzed known crossover hotspots (*25, 27*). The H4K8la map exhibited a strong correlation with both Crick and Watson sites of DMC1-SSDS hotspots (*26*), and enrichment of SPO11-oligo was observed near DMC1-SSDS hotspot centers (**Figures 3F)**. Additionally, our data show that H4K8la peaks are highly correlated with RAD51 and RPA2 sites (**Figures 3F)**. H4K8la peaks also show correlation with known crossover hotspots (**Figures 3F**; **Supplementary Figure 5**).

Recent studies have shown that the presence of H3K4me3 is necessary but not sufficient for DSB formation (*21, 28*). Despite its ubiquity at DSB sites, H3K4me3 can also be found at other functional locations, such as gene promoters. Emerging evidence suggests additional histone marks, particularly in histones H3 and H4 (e.g., H3K9ac, H3K27ac, H3K4me1), may also play a pivotal role at these sites (*21, 24*). Taken together, these data suggest that additional histone marks are required for a comprehensive description of DSB sites (**Supplementary Figure 6A and 6B**). A moderate depletion at DSB hotspots was observed for H4K8la at the leptotene stage compared to H3K4me3 (**Supplementary Figure 6B**). Interestingly, H4K8la and H3K27ac marks that overlapped with SPO11-oligo hotspots were formed during zygotene, mirroring the behavior of H3K4me3 at zygotene. This underscores the potential influence of other chromatin characteristics on meiotic recombination.

### H4K8la, but not histone H4 K5K8K12 acetylation, is associated with recombination hotspots

In addition to lactylation, histone H4 K8 can undergo acetylation (H4K8ac). To ascertain whether H4K8la is specifically involved in recombination hotspots, we compared H4K8la with H4K8ac, as well as histone H4 acetylation at adjacent lysine sites, including H4K5ac and H4K12ac (*21*). Consistent with a previous study indicating that H4K8ac, H4K5ac, and H4K12ac are not detected at recombination hotspots (*21*), our data reveal that, while the H4K8la signal at SPO11-oligo- and DMC1-SSDS hotspots during zygotene is robust, the signals for H4K8ac, H4K5ac, and H4K12ac at these sites are weak **Figure 4A - 4C; Supplementary Figure 7**).

**Figure 4.**
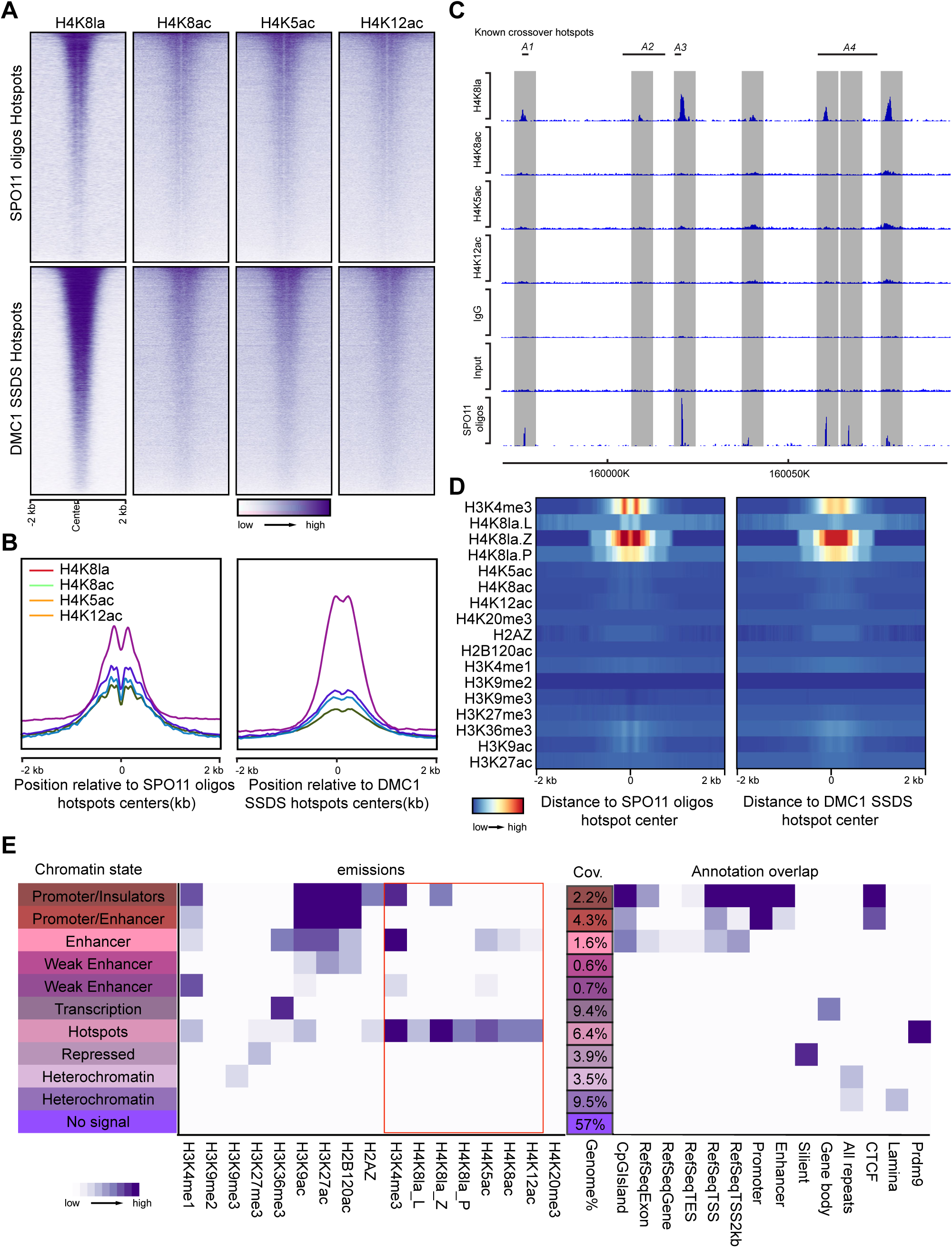
Comparison of H4K8la with other histone H4 modifications at meiotic recombination hotspots. **(A)** Heatmaps showing the coverage of H4K8la, H4K8ac, H4K5ac and H4K12ac in both SPO11 oligos and DMC1 SSDS hotspots in zygotene. Each row in these heatmaps corresponds to one hotspot. These hotspots are organized in descending order based on the abundance of SPO11 oligos and the intensity of DMC1 SSDS ChIP-seq peaks. **(B)** Distribution of H4K8la, H4K8ac, H4K5ac and H4K12ac around the centers of SPO11 oligos and DMC1 SSDS hotspots. **(C)** Genome browser view of SPO11oligos and ChIP-seq of H4K8la, H4K8ac, H4K5ac and H4K12ac at four known crossover hotspots (*A1–A4*). **(D)** Enrichment of the indicated histone marks at SPO11 oligos and DMC1 SSDS hotspots. **(E)** Heatmap of the emission parameters for a 11-state ChromHMM model based on ChIP-seq results of 14 histone modifications.

We proceeded to perform an integrated analysis of H4K8la with 14 histone methylation and acetylation marks obtained from published datasets (**Figure 4D**) (*29, 30*). Several of these histone marks have been previously identified at recombination hotspot sites in mice (*21*) or reported to exhibit enrichment at double-strand break (DSB) sites in other organisms (*31*). Our findings indicate that, similar to H3K4me3, H4K8la is strongly associated with recombination hotspots marked by SPO11-oligos and DMC1-SSDS.

We conducted a comprehensive integrated analysis of H4K8la with these 14 histone methylation and acetylation marks using ChromHMM, a hidden Markov model-based tool (*32*). This allowed us to identify co-occupancy patterns of histone modifications defining distinct chromatin "states" (*33*). Utilizing both new and previously published nucleosome-resolution ChIP-seq data related to histone modifications or variants (*29, 30*), we established an 11-state model. Our analysis revealed multiple chromatin states, encompassing "Promoter and insulator regions," "Enhancer elements," "Transcriptionally active regions," "Polycomb repressive complex-repressed domains," "Heterochromatic zones," and notably, "Hotspots" (**Figure 4E**). Remarkably, among these chromatin states, only "Hotspots" exhibited robust enrichment for PRDM9-binding sites and was characterized by a distinct collection of histone modifications, including H3K4me3 and H4K8la (**Figure 4E**). In contrast to other histone acetylation marks typically found at enhancers, such as H3K27ac and H2BK120ac, we observed that CTCF signal was enriched at promoter and insulator regions. Interestingly, H4K8la at zygotene was also enriched at promoter and insulator regions, suggesting the involvement of H4K8la in 3D genome organization (**Figure 4E**).

### H4K8la correlates with dynamic 3D genome organization

Recent studies have contributed multiple meiotic Hi-C datasets for spermatocytes (*4, 34*). We combined datasets to analyze the chromatin structure at the leptotene, zygotene, and pachytene stages during male germ cell meiosis alongside our H4K8la sites. Other chromatin states, including occupancy patterns of CTCF, meiotic-specific cohesin (RAD21L, REC8), and double-strand breaks (DSBs) (SPO11-oligos and DMC1-SSDS), were also considered. All datasets were uniformly mapped to a consistent set of 10-kb bins across the autosomal chromosomes (**Fig. 5A**). Interestingly, we observed that, in addition to DSB hotspots, H4K8la sites also strongly correlated with meiosis-specific cohesion sites where RAD21L and REC8 are located (**Fig. 5B**). These meiosis-specific cohesion sites do not overlap with DSBs hotspots (**Fig. 5B**). This demonstrates that the H4K8la plays two separated roles in meiotic prophase I.

**Figure 5.**
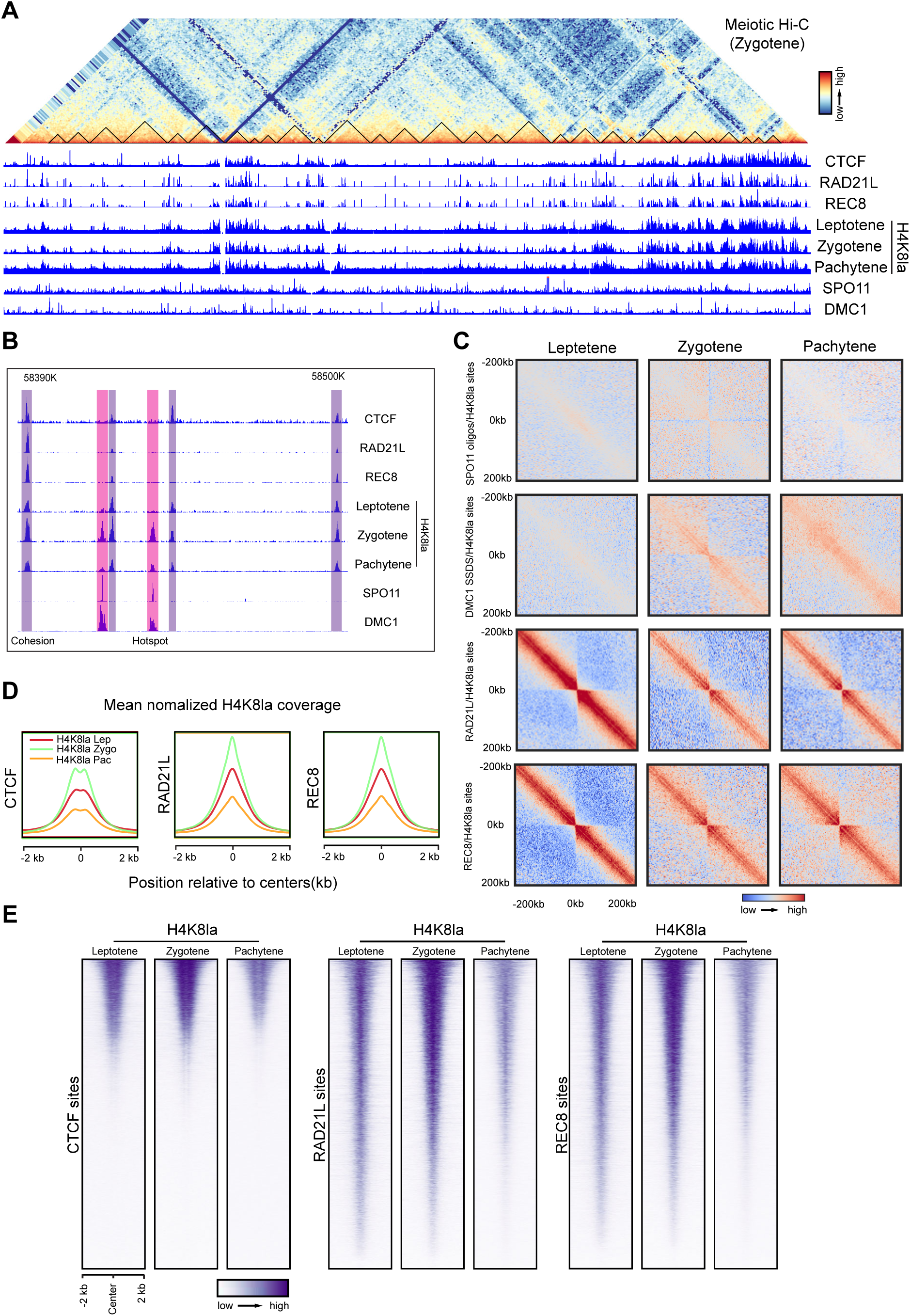
Histone lactylation dynamics at meiotic specific cohesion sites. **(A)** Genome browser view of a representative region on mouse mm10 Chromosome 4. Zygotene Hi-C contact map is shown as a heatmap with ChIP-seq tracks of cohesin subunits, RAD21L, REC8 and CTCF, DSB hotspot tracks of SPO11 oligos and DMC1 SSDS, and ChIP-seq for H4K8la. **(B)** CTCF, RAD21L, REC8, SPO11, DMC1 and H4K8la map in zoom-in window around DSB hotspot sites and cohesion sites. **(C)** Normalized chromatin contact matrices, symmetric-averaged across H4K8la overlapped with SPO11 Oligos, DMC1 SSDS ChIP-seq, RAD21L and REC8 sites in leptotene, zygotene and pachytene Hi-C datasets. **(D)** Distribution of H4K8la coverage around centers of CTCF, RAD21L and REC8 sites. **(E)** Heatmaps showing the coverage of H4K8la reads in CTCF, RAD21L and REC8 site. Each row in these heatmaps corresponds to one of site.

We conducted a further analysis of the positioning and distribution of chromatin loops by generating composite pileups of interactions among co-occupied sites involving H4K8la with SPO11-oligo (8,167), DMC1-SSDS (9,457), RAD21L (10,479), and REC8 (10,541) within a ±200-kb region. Our data reveal a distinct induction of interactions between double-strand break (DSB) sites and H4K8la sites in meiotic Hi-C maps, peaking in zygotene when DSBs are predominantly introduced and diminishing in pachytene when DSBs are repaired (**Fig. 5C**). In contrast, RAD21L/H4K8la and REC8/H4K8la co-occupied sites exhibit the strongest enrichment of interactions in leptotene, as evidenced by the prominent central signals in the pileup heatmaps (**Fig. 5C**).

To further explore the correlation between H4K8la and cohesion activity, we mapped CTCF, RAD21L and REC8 ChIP-Seq binding sites in H4K8la during leptotene to pachytene, suggesting potential dynamic shift from leptotene to pachytene (**Fig. 5D and E**). Our heatmap analysis revealed a strong correlation between REC8 and RAD21L with H4K8la, although CTCF did not exhibit a strong correlation. The signal profiles from CTCF, RAD21L, and REC8 ChIP-Seq binding sites were consistent with the heatmap analysis, suggesting a dynamic cohesion program in meiotic prophase.

## Discussion

Our data unveil, for the first time, the involvement of histone lysine modifications in meiosis. As expected, H4K8la is associated with the gene activation required for this process. However, intriguingly, our findings indicate that H4K8la is not only detected at the recombination hotspots but also at the cohesion sites flanking these hotspots. The observed increase in H4K8la modification from leptotene to zygotene suggests the generation of new H4K8la modifications in proximity to the original ones, implying a double-strand break (DSB)-dependent mechanism for H4K8la modification. Conversely, the decrease in H4K8la signal from zygotene to pachytene may indicate the progression of DSB repair. Overall, our results reveal a precise spatiotemporal regulation of H4K8la dynamics associated with recombination hotspots during the leptotene to pachytene stages. In our analysis, both the SPO11-oligo and DMC1-SSDS peaks precisely overlapped with a trough situated between fluctuating H4K8la peaks.

To further investigate, mice with a genetic deletion of machinery in the meiotic recombination pathway would be valuable for dissecting the role of H4K8la. For example, if H4K8la is not present at meiotic recombination hotspots in Spo11-deficient germ cells, this data would support the idea that H4K8la at recombination hotspots depends on meiotic double-strand break (DSB) formation and that meiotic DSB formation precedes H4K8la. Additional experiments utilizing Spata22- and Rpa2-deficient germ cells could determine whether H4K8la at recombination hotspots depends on single-strand break resection. If so, Dmc1-knockout germ cells could be used to test whether H4K8la at recombination hotspots depends on strand invasion. Conversely, if H4K8la remains at meiotic recombination hotspots in Spo11-deficient germ cells, this data would suggest that H4K8la at recombination hotspots precedes DSB formation. Prdm9-knockout germ cells could then be employed to test whether H4K8la at recombination hotspots depends on H3K4me3 at these sites. These experiments are currently in process.

## Acknowledgement

We thank Clark Bloomer, Rosanne Skinner, Veronica Cloud, and Yafen Niu at the University of Kansas Medical Center Genomics Core, which is supported by Kansas Intellectual and Developmental Disabilities Research Center (NIH U54 HD 090216), the Molecular Regulation of Cell Development and Differentiation—COBRE (P30 GM122731-03)—the NIH S10 High-End Instrumentation Grant (NIH S10OD021743) and the Frontiers CTSA grant (UL1TR002366). This work was supported by the National Institutes of Health (NIH) grant, R01HD-103888, and the Department of Cell Biology and Physiology to N.W.. X.Z. is a recipient of KUMC Research Institute Lied Pilot Award. X.Z. is a recipient of Kansas IDeA Network of Biomedical Research Excellence (K-INBRE) Developmental Research Project Program Award from the National Institute of General Medical Sciences of the National Institutes of Health under grant number P20 GM103418. The content is solely the responsibility of the authors and does not necessarily represent the official views of the above-mentioned funders.

## Materials and Methods

### Animals

All experiments involving animals were conducted in accordance with the approved protocol by the Institutional Animal Care and Use Committee (IACUC) at the University of Kansas Medical Center, strictly adhering to its regulatory and ethical guidelines. The animals were housed in a specified pathogen-free facility with a 12-hour light/dark cycle, and they had ad libitum access to food and water.

### Antibodies

Pan anti-Kla (PTM-1401), anti-H2BK15la (PTM-1426RM), anti-H3K18la (PTM-1427RM), anti-H3K14la (PTM-1414RM), anti-H2BK16la (PTM-1424RM), anti-H2AZK11la (PTM-1422RM), anti-H4K8la (PTM-1415RM), anti-H4K12la (PTM-1411RM), anti-H3K9la (PTM-1419RM), anti-H4K16la (PTM-1417RM), anti-H4K5la (PTM-1407RM) and anti-H4K8la (PTM-1415) antibodies were purchased from PTM Bio Inc.. Anti-H3K4me3 (ab8580) was purchased from Abcam. Anti-SYCP3 (sc-74569) was purchased from Santa Cruz. Anti-phospho-Histone H2A.X (Ser139) (05-636-I) was purchased from Millipore.

### Histological analysis

The dissected mouse testis tissues underwent fixation using a 4% paraformaldehyde (PFA) solution at 4°C overnight. Subsequently, the fixed tissues were embedded in paraffin and sectioned into slices with a thickness of 6 μm.

### Immunofluorescence staining

The sections of mice testis tissue underwent deparaffinization with xylene, followed by a sequential rehydration process using ethanol concentrations of 95%, 80%, and 70%. Antigen retrieval was achieved by heating the sections in a 10 mM citrate buffer (pH 6.0) using a microwave. Subsequent washing was performed using Phosphate-Buffered Saline (PBS) with 0.05% Tween 20. This step was followed by an overnight incubation with primary antibodies at 4°C and then with secondary antibodies at room temperature for one hour. Images were captured using a Nikon A1R confocal microscope and subsequently processed with NIS-Elements Viewer v5.21 and Adobe Photoshop software.

### Chromosome spreads

The testes were dissociated after removing the tunica albuginea in 15 mL tubes containing 5 mL TIM buffer (104 mM NaCl, 45 mM KCl, 1.2 mM MgSO4, 0.6 mM KH2PO4, 6.0 mM sodium lactate, 1.0 mM sodium pyruvate, and 0.1% glucose) with 1 mg/ml collagenase IV. The tissue was left shaking for 1 h at room temperature. After incubation, the TIM buffer was removed following centrifugation for 5 min at 300 × g at room temperature. A 1 ml tip was used to disperse the tissue further by pipetting up and down for 5 min. The single-cell suspension was resuspended in 500 µl TIM. A 75 mM sucrose solution was added to the tube. Then, 200 µl of the cell mix was transferred to Superfrost plus glass slides (Fisher Scientific) coated with a thin layer of 1% paraformaldehyde (PFA). The slides were half-dried in a closed slide box for 1 h and then dried with a half-open lid for 2 h at room temperature. Subsequently, the slides were washed with 0.4% PhotoFlow (Nikon), air-dried, and stored at −80 °C.

### Synchronization of spermatogenesis

Synchronization of spermatogenesis was accomplished based on a published protocol (*22*). In brief, male mice received daily subcutaneous injections of WIN18,446 (0.1 mg/g body weight, Santa Cruz Biotechnology) from postnatal day 2 to 8 and a single dose of retinoic acid (RA) (0.0125 μg/g body weight, MilliporeSigma) on day 9. The animals were sacrificed approximately 8, 10, and 12 days post-RA treatment to harvest testes, which were enriched for leptotene, zygotene, and pachytene cells.

### Chromatin immunoprecipitation (ChIP) and data analysis

ChIP was performed on testicular cells, which were dissociated after tunica albuginea removal and fixed in 1% formaldehyde for 10 minutes. Quenching was carried out with homogenized tissue washed twice in PBS. Cell lysis took place in 1 ml of EZ-ChIP kit lysis buffer (Millipore, 17-371RF), and chromatin was sheared into approximately 1000 bp fragments via sonication using a Diagenode Pico Bioruptor. The sample underwent pre-clearing with Protein G beads, and a 10 µl aliquot of this pre-cleared chromatin served as the ’input’ sample. The remaining chromatin was incubated with either H4K8la antibody or preimmune IgG overnight at 4°C, followed by a 2-hour incubation with Protein A/G Agarose beads. Subsequent washing steps preceded chromatin elution from the beads with 1% SDS and 0.1M NaHCO3 (pH 9.0) at 65°C, with crosslink reversal occurring overnight at the same temperature. DNA deproteinization was conducted at 45°C for 2 hours, followed by purification. Enrichment was assessed through sequencing.

The processing of ChIP-seq data began with the removal of Illumina adapter sequences from paired-end reads using Cutadapt (v2.10). Reads were selected with a minimum length of 25 bp. Alignment to the mouse mm10 reference genome was conducted with Bowtie2 (v2.4.2) using local, very sensitive settings and specified argument ranges.

After alignment, duplicate reads were identified using the Picard MarkDuplicates tool, and BAM files were filtered via SAMtools to exclude unmapped reads, non-primary alignments, failed quality checks, and duplicates (-F 1804 -f 2).

Peak calling was performed using MACS2 with a q-value threshold of 0.05. Consensus peak sets for biological replicates of each histone mark or factor were obtained by intersecting peak sets using BEDTools intersect, retaining peaks present in both replicates. Visualization of the data included profile plots generated by Deeptools. For genome browser visualization, normalized BigWig files were generated using DeepTools bamCoverage, with parameters set for bin size, processor count, normalization, and genome size.

### Author contributions

X.Z. conceptualized the project and performed all the experiments with helps from Y.L.. X.Z. and N.W. designed the experiments, analyzed data, and wrote the manuscript.

