## Supplementary Materials for "Multifaceted Roles of Histone Lysine Lactylation in Meiotic Gene Dynamics and Recombination"

The PDF file includes:

Figs. S1 to S8

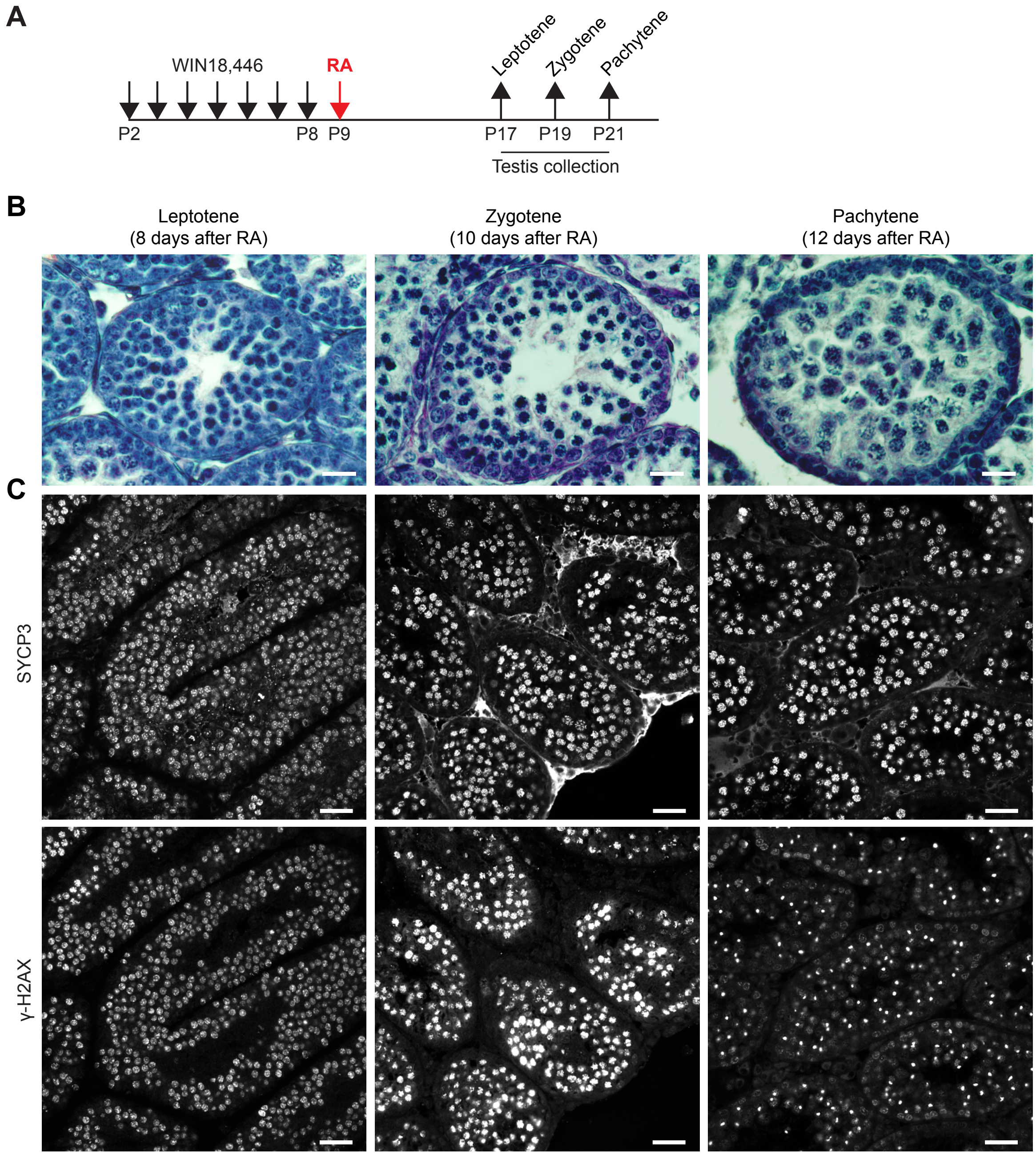


**Supplementary Figure 1**

**Supplementary Figure 1. Synchronized spermatogenesis.**

**(A)** Schematic of the synchronization of spermatogenesis using WIN18,446 and retinoic acid (RA), to enrich meiotic stage-specific testicular germ cells.

**(B)** Histology of testes from mice with synchronized spermatogenesis. Scale bar, 25 μm.

**(C)** Immunofluorescence (IF) staining of SYCP3 and γH2AX in testicular cross sections from male mice with synchronized spermatogenesis showed a synchronized stages of meiosis (leptotene, zygotene, and pachytene). Scale bar, 50 μm.

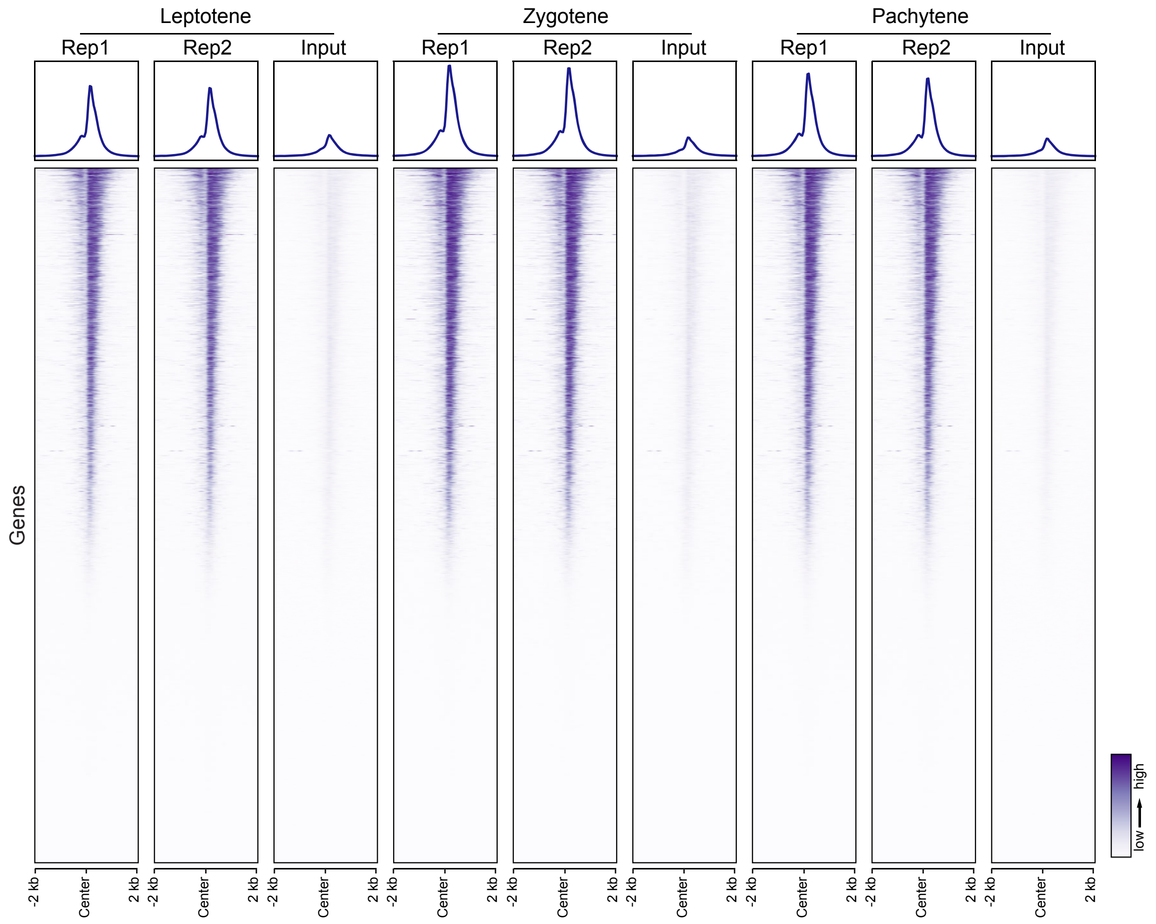


**Supplementary Figure 2**

**Supplementary Figure 2. Biological replicates of H4K8la ChIP-seq.**

Biological replicates of H4K8la ChIP-seq from leptotene to pachytene.

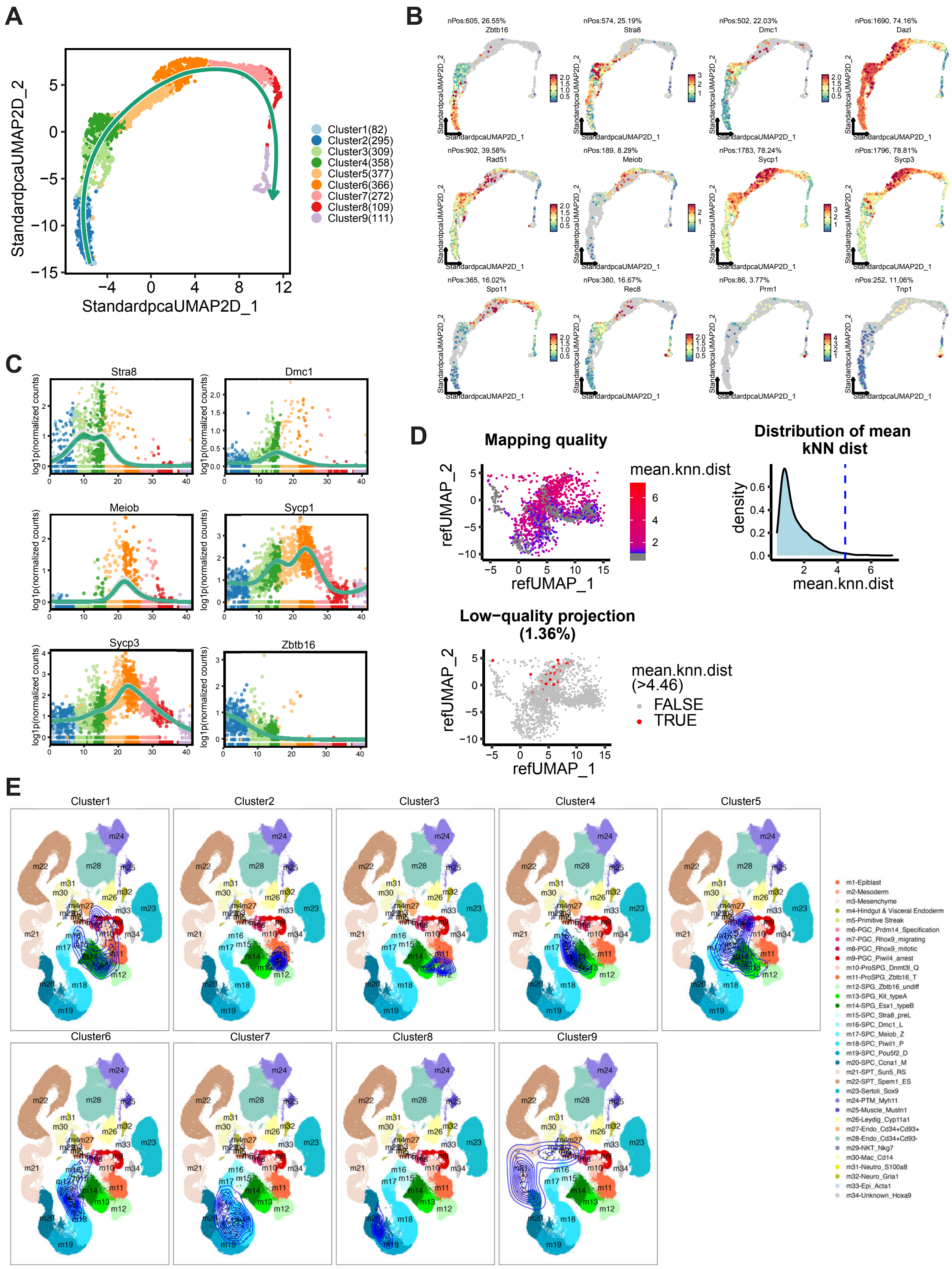


**Supplementary Figure 3**

**Supplementary Figure 3. scRNA-seq analysis of testicular germ cells.**

**(A)** Uniform manifold approximation and projection (UMAP) plot along the projected pseudo-time of cells captured from mouse testicular germ cells colored by clusters.

**(B)** UMAP plots of distribution of known marker genes in each cluster.

**(C)** Expression levels of marker genes shown in pseudo-time.

**(D)** The mapping quality of the projected cells. Dots are colored by mean KNN distance (mean.knn.dist). Distribution of mean.knn.dist corresponding to panel (D). The dashed line indicating the cutoff value (mean.knn.dist > 4.46) represents low-quality projected cells on embeddings.

**(E)** Reference of the mouse testicular cells from 6.5 days post coitum (dpc) to adults. Our datasets consisting of biological replicates were mapped to the testis reference by using ProjectSVR package. 2D density plots show the projected cells onto the mouse testicular cell reference, which confirms that the cells are at leptotene (Cluster 6), zygotene (Cluster 7), and pachytene (Cluster 8) stages.

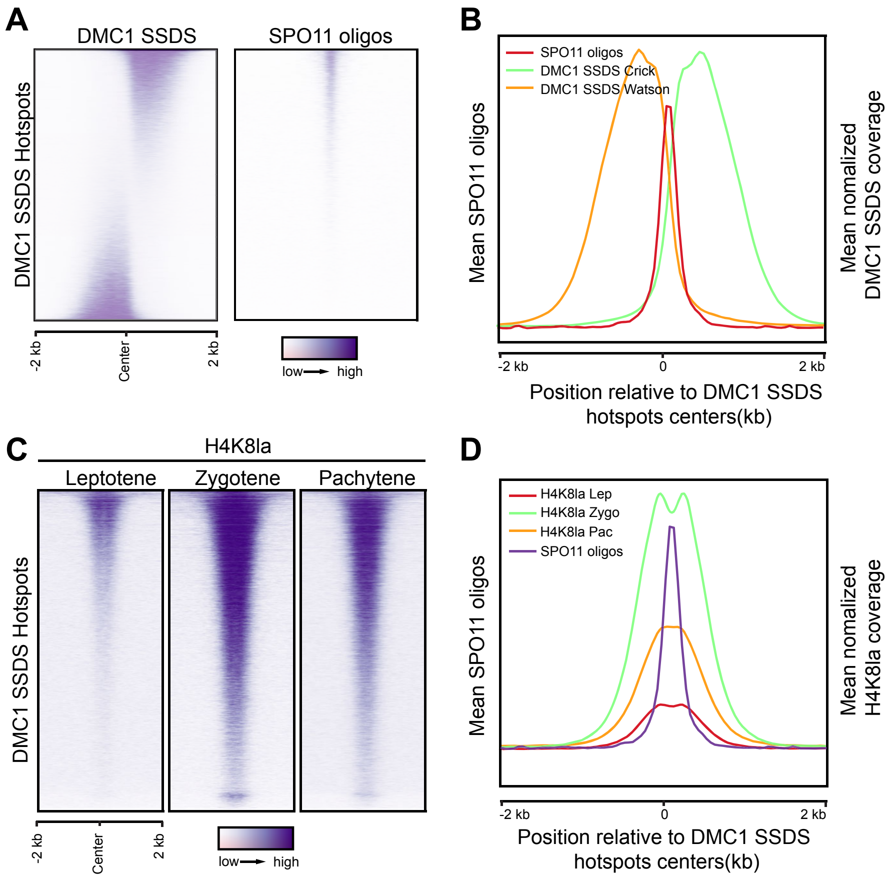


**Supplementary Figure 4**

**Supplementary Figure 4. Histone lactylation dynamics at SSDS hotspots**

**(A)** Heatmaps showing the coverage of DMC1 SSDS ChIP-seq reads and SPO11 oligos in hotspots identified in the C57Bl/6 strain. Each row in these heatmaps corresponds to one hotspot. These hotspots are organized in descending order based on the abundance of DMC1 SSDS coverage.

**(B)** Distribution of both the Crick and Watson strands of DMC1 SSDS coverage and SPO11 oligos around the centers of the DMC1 SSDS hotspots.

**(C)** Heatmaps showing the dynamic H4K8la signals at DMC1 SSDS hotspots. Each row in these heatmaps represents a specific hotspot-associated peak, extending ±2 kb from the peak center. These peaks are organized in descending order, based on their DMC1 SSDS density.

**(D)** Distribution of SPO11 oligos and hotspots associated H4K8la peak coverage around centers of DMC1 SSDS hotspots.

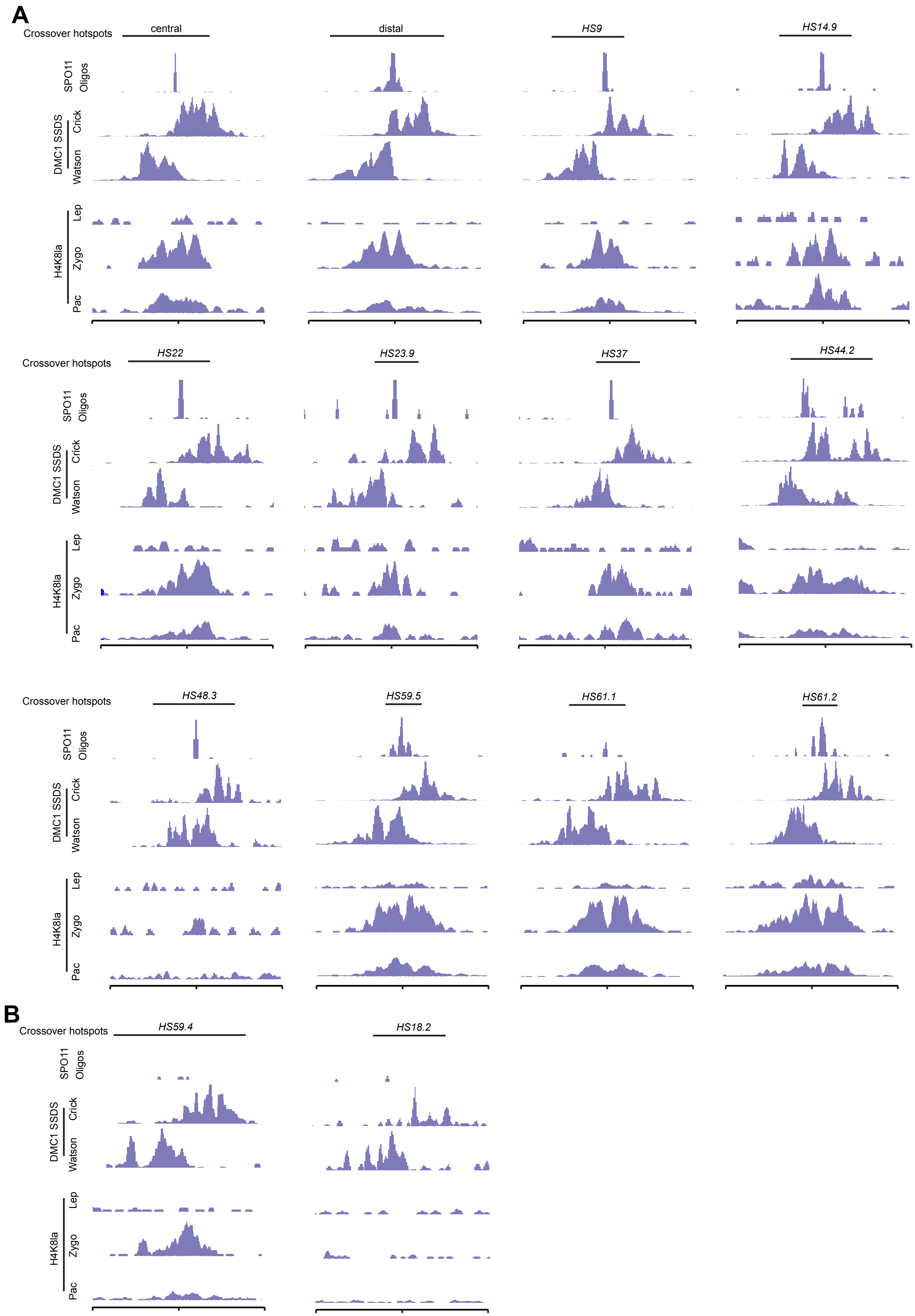


### Supplementary Figure 5

**Supplementary Figure 5. Known crossover hotspots contain hotspot associated H4K8la that are present from leptotene to pachytene.**

**(A)** Genome browser view of H4K8la reads coverage on known crossover hotspots with both SPO11 oligo and DMC1 SSDS signals in different stages of leptotene to pachytene. The positions of crossover hotspots are highlighted.

**(B)** Genome browser view of H4K8la reads coverage on known crossover hotspots with only DMC1 SSDS signals in different stages of leptotene to pachytene. The positions of crossover hotspots are highlighted.

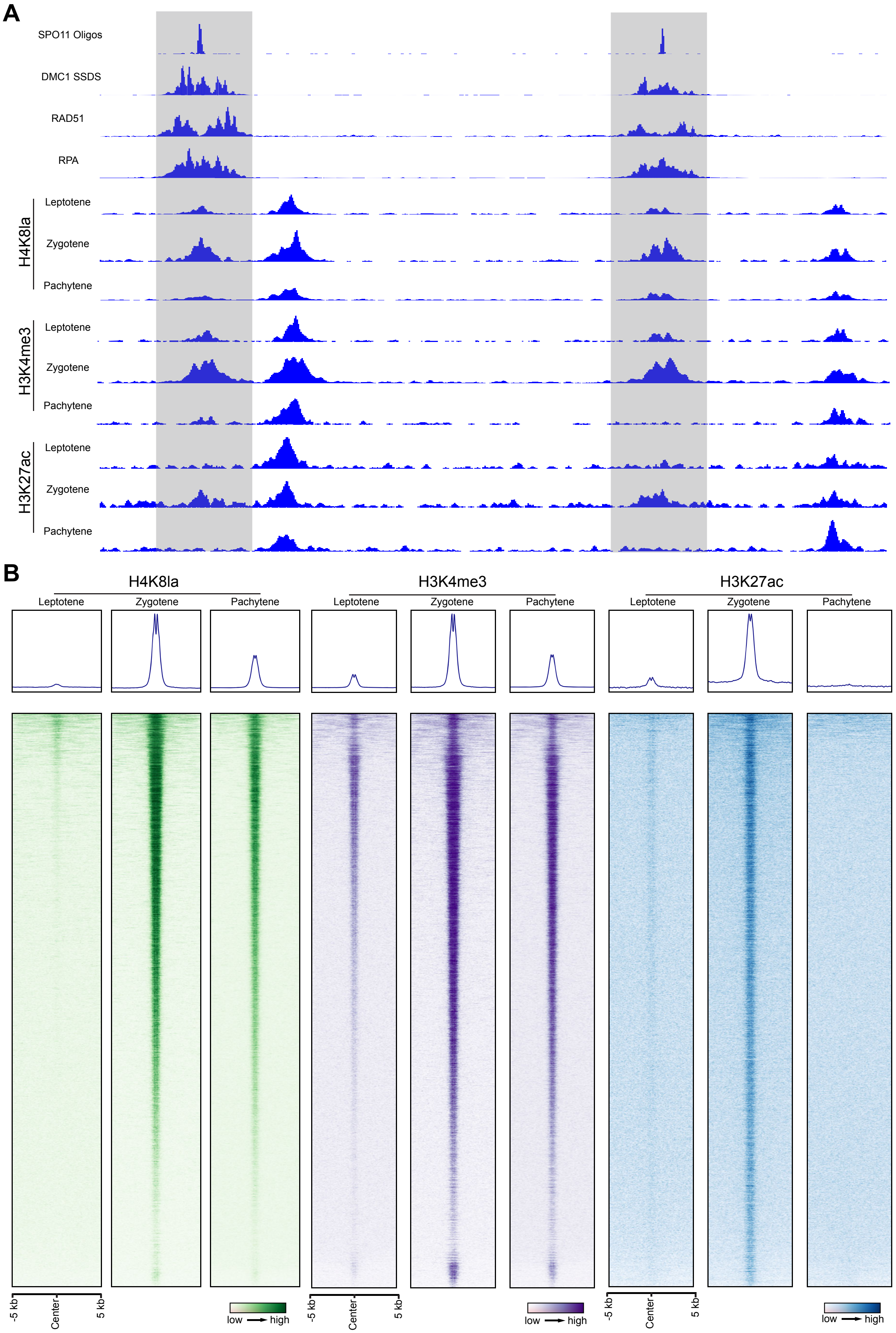


### Supplementary Figure 6

**Supplementary Figure 6. Various histone marks at recombination hotspots.**

**(A)** An integrated genome track of SPO11 oligos, DMC1 SSDS, ChIP-seqs for RAD51 and RPA, H4K8la ChIP-seq, H3K4me3 ChIP-seq, and H3K27ac ChIP-seq reads in a segment of chromosome 1 with two recombination hotspots.

**(B)** Heatmaps of H4K8la (left), H3K4me3 (center) and H3K27ac (right) on SPO11 oligo hotspot associated H3K4me3 peaks from leptotene to pachytene stages. Each row represents a SPO11 oligo hotspot peak of ±5 kb around the peak center and ranked based on SPO11-oligo.

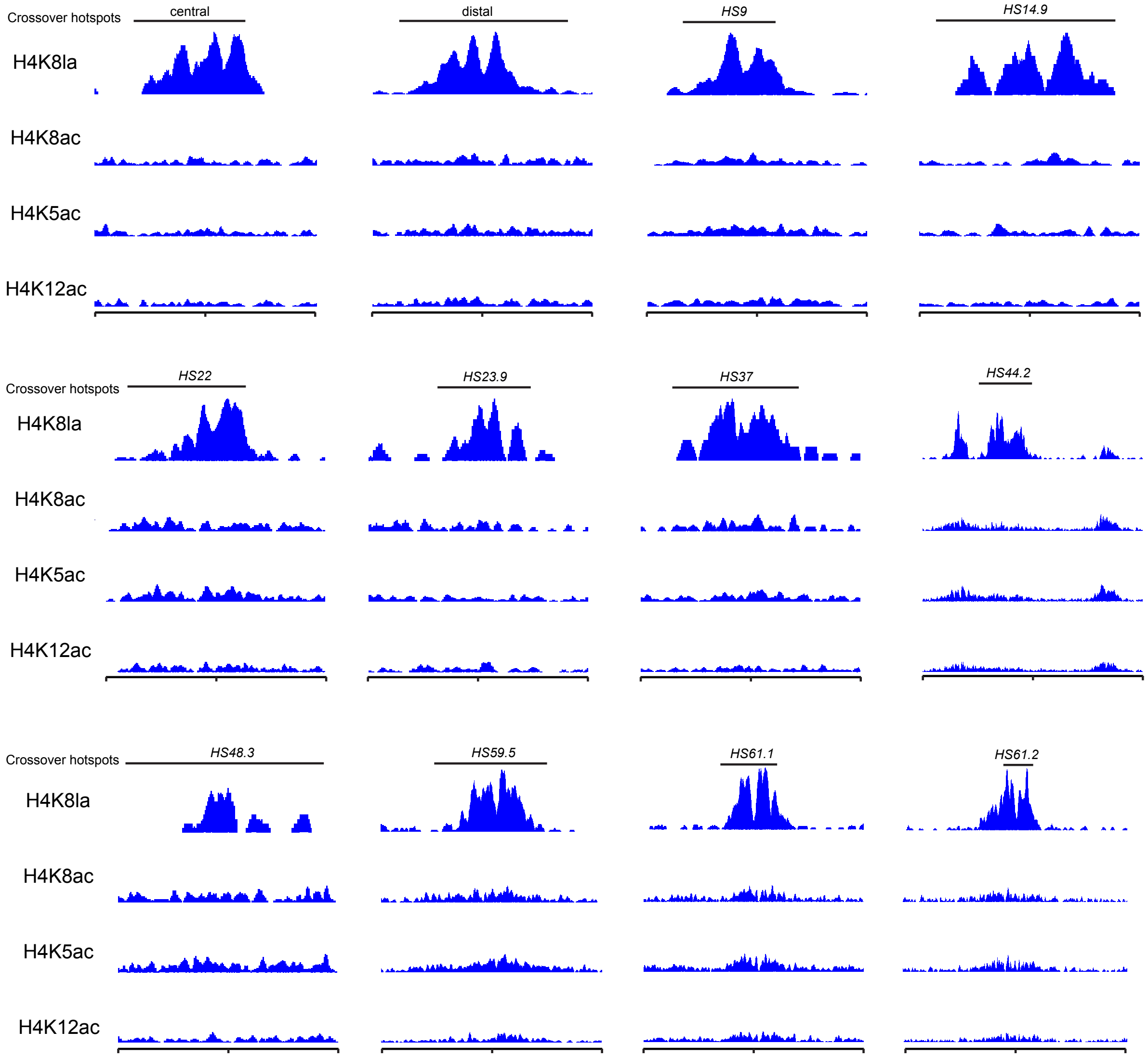


**Supplementary Figure 7**

**Supplementary Figure 7. Comparison of H4K8la with H4K5ac, H4K8ac, H4K12ac at known crossover hotspots at zygotene.**

Genome browser view of H4K8la, H4K5ac, H4K8ac, and H4K12ac coverage at known crossover hotspots at zygotene. The positions of crossover hotspots are indicated.

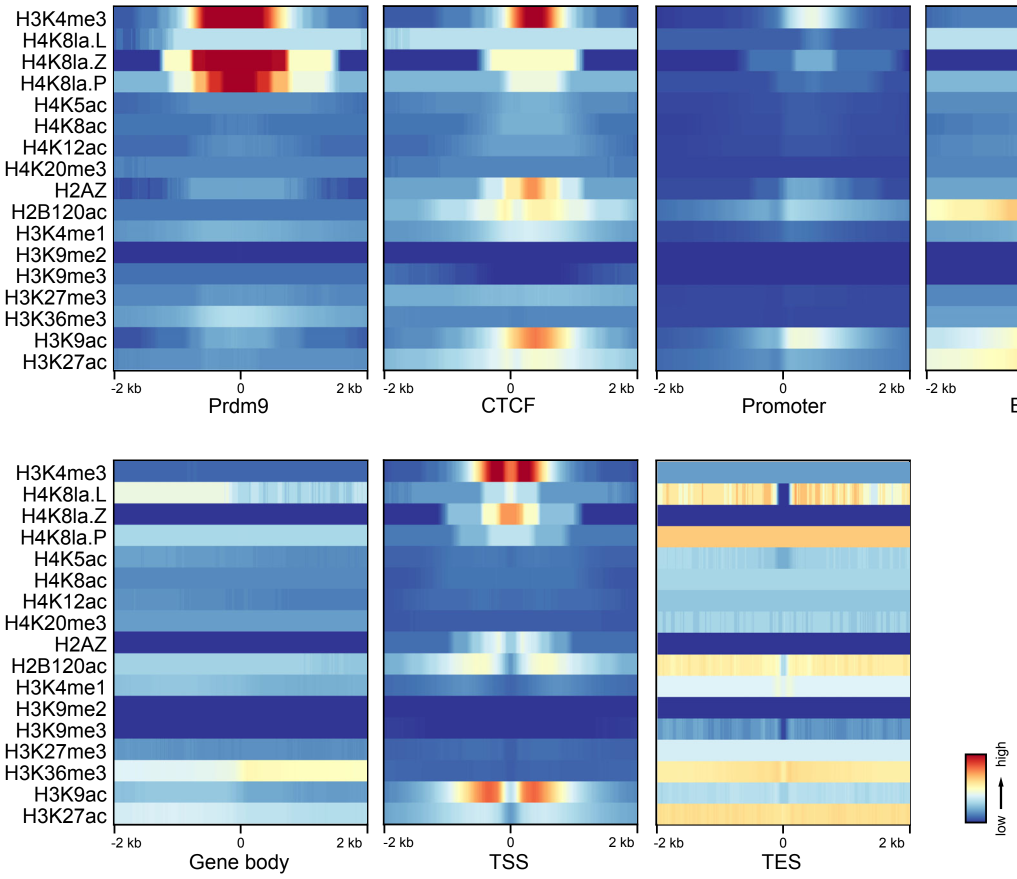


### Supplementary Figure 8

**Supplementary Figure 8. Enrichment of histone modifications at functional genomic elements.**

Coverages of 14 histone modification marks were enriched around genomic functional sites. Genomic functional sites data were sourced from 11 collected chromatin states.
